# Readiness to retrieve memories of different ages: An ERP dissociation of orienting and difficulty

**DOI:** 10.1101/2022.07.25.501431

**Authors:** Emily K. Leiker, Jeffrey D. Johnson

## Abstract

A well-established finding in episodic memory research is that the likelihood of successful retrieval is affected by the ‘age’ of the targeted memory (i.e. based on the duration of the encoding-retrieval interval). Although such findings are usually interpreted in terms of the memory trace changing in quality or accessibility over time, there is growing evidence that memory performance also depends on preparatory attention and strategic orienting at the time of retrieval. Here, we investigated such readiness to retrieve, as indexed by event-related brain potentials (ERPs), when memory retrieval was aimed toward candidate traces of different ages. As retrieving remote versus recent memories is associated with increased difficulty, the current study was designed to dissociate difficulty from the memory age effects. Subjects encoded two lists of pictures separated by one week, with the items in each list presented once or four times. A series of memory tests targeting pictures from only one week-by-repetition condition at a time were then completed, while new pictures and those from the other conditions were rejected. Consistent with previous results, new-item ERPs were more positive over posterior scalp when recent compared to remote lists were targeted. Importantly, this difference remained when difficulty was matched, and the ERP correlates of difficulty were distinct in both timing and polarity from those of orienting. Together, these findings provide further support for the idea that the age of a targeted memory alone places different demands on readiness and can lead to the differential adoption of retrieval orienting strategies.

## 1. INTRODUCTION

A long-standing empirical observation about human memory is that successful retrieval becomes less likely as more time passes since the encoding episode (for reviews, see Roediger et al., 2010; Wixted, 2004). To date, the research on such forgetting effects has largely focused on either characterizing the neurocognitive at encoding that result in more durable memory traces (e.g., Craik & Lockhart, 1972; Paller & Wagner, 2002; Sneve et al., 2015; Uncapher & Rugg, 2005; Wagner et al., 1998) or understanding how factors such as consolidation, decay, and interference change memory traces over time (Dewar et al., 2007; Dudai, 2004; Frankland & Bontempi, 2005; Wixted, 2005). There has been little research, though, on how strategic processes at the time of retrieval might be differentially engaged depending on when a memory was encoded and whether there are consequences of such strategies on retrieval success. The current study sought to further characterize the functional nature of retrieval strategies targeting the age of a memory and to dissociate them from another prominent difference related to memory age: retrieval difficulty.

One form of strategic processing that has been distinguished from effects that directly correspond to memory outcome (i.e. success) is retrieval orienting (or orientation). Orienting is thought to constrain the retrieval search by processing the cue in a manner aligned with the target memory (e.g., Herron & Rugg, 2003a; Jacoby et al., 2005; Robb & Rugg, 2002), thereby increasing cue-target overlap and improving likelihood of retrieval success (see Morris, Bransford, & Franks, 1977; Rugg et al., 2008; Rugg & Wilding, 2000; Tulving & Thomson, 1973). Evidence for the adoption of different retrieval orientations comes predominantly from studies employing event-related potentials (ERPs) while memories encoded under different conditions are targeted in separate retrieval blocks, thus promoting the sustained engagement of distinct cognitive sets (e.g., Herron & Wilding, 2005; Johnson et al., 1997; Johnson & Rugg, 2006; Robb & Rugg, 2002; Rosburg et al., 2013; Rugg et al., 2000; Wilding, 1999). Additionally, these studies typically restrict analysis to ERPs elicited by correctly-rejected new (non-studied) test items (cf. studies that analyze pre-stimulus effects across all test items; e.g., Herron & Evans, 2018; Herron & Wilding, 2004, 2006), thus avoiding potential confounds due to the rate of retrieval success or engagement of post-retrieval processes across different targeted memory types. In one example study taking this approach, Robb and Rugg (2002) identified a retrieval orientation effect beginning by around 500 ms post-stimulus onset, with ERPs exhibiting more positivity when targeting previously-encoded words compared to pictures. Importantly, these ERP effects were evident for cues that were identical in form (words in both cases) across the two conditions and were not studied, thus ruling out any differences due to perception, retrieval success, and post-retrieval monitoring. There is also increasing evidence that differences in memory performance associated with healthy aging may depend in part on the ability to engage such strategic retrieval processes (for reviews, see Madore & Wagner, 2022; Morcom, 2016; Wilckens et al., 2012).

The approach of studying retrieval orientation described above has also recently been adapted to investigate strategic orienting to memory age. In an initial study, Roberts, Tsivilis, and Mayes (2014) investigated retrieval orientation in the context of pictures that were encoded either 48 hours or 40 minutes earlier. Topographically widespread ERP differences were evident when targeting items from only a single encoding period at a time compared to items from both encoding periods simultaneously. Although the ERPs did not distinguish between which encoding period was targeted, these findings suggested that subjects can use memory age, albeit in a somewhat general manner, to orient retrieval. In a follow-up study, Johnson and McGhee (2015) sought to test for differential orienting to specific memory ages by expanding the encoding periods from 30 minutes to 1 week earlier. A key feature of this study was that it employed an exclusion procedure, in which subjects are instructed to respond positively to studied items from a particular condition (e.g., 1 week earlier), while rejecting new items as well as those studied in other conditions (e.g., 30 minutes ago; Jacoby, 1991). The exclusion task presumably involves adopting a strategy of orienting retrieval toward the targeted class of items (Herron & Rugg, 2003b; Herron & Wilding, 2005; Wilding & Rugg, 1997), and accordingly, larger ERP orientation effects have been demonstrated when memory is tested under exclusion as opposed to simple recognition procedures (Johnson & Rugg, 2006). Johnson and McGhee (2015) observed more positive-going ERPs over posterior scalp when recent (30 minutes earlier) compared to remote (one week ago) items were targeted in distinct retrieval blocks, providing novel evidence that individuals can adjust their orienting strategy to the specific memory age of targets.

One consequence of using longer delays between encoding periods is that the adoption of orienting strategies can be confounded with differences in difficulty. Whereas Johnson and McGhee (2015) attempted to reduce such effects in a post hoc manner by matching trials according to response time (RT) and splitting subjects into groups that showed larger versus smaller difficulty differences (also see Dzulkifli et al., 2004), an alternative method is to dissociate the effects via direct manipulation that matches difficulty across orienting conditions. Here, we employed an experimental design that was similar to but expanded upon that of Johnson and McGhee (2015). Subjects completed two encoding phases separated by one week, with half of the items in each phase presented once and the other half presented four times. The memory retrieval phase then used an exclusion task procedure in which each combination of items from the remote/recent and repetition conditions was targeted in separate blocks. The key contrast involved ERPs for correctly-rejected new items in the test block targeting remote items presented four times and those in the block targeting recent items studies once. Remaining orientation-related ERP differences in the face of matched retrieval difficulty, as well as evidence that orientation and difficulty can be dissociated on the basis of scalp topography or polarity (for a similar approach, see Robb & Rugg, 2002), would provide further support that subjects can orient retrieval toward memory age.

## 2. METHODS

The materials, data, and analysis code used for this study are available at https://osf.io/j8qm3/. No part of the study procedures or analyses was pre-registered prior to the research being conducted. We report how we determined our sample size, all data exclusions, all inclusion/exclusion criteria, whether inclusion/exclusion criteria were established prior to data analysis, all manipulations, and all measures in the study.

### 2.1 Subjects

Twenty-four undergraduates from the University of Missouri (MU) participated for course credit. Our target sample size was based on those of prior ERP studies of retrieval orienting instead of an *a priori* power analysis. Based on the primary effect of interest in our previous study of orienting to memory age – the *F*-test for the 500-800 ms orienting difference for new items in Johnson and McGhee (2015) – we conducted a *post hoc* power analysis. The analysis indicated that a sample size of 15 subjects would have given an 80% chance of rejecting the null hypothesis for this large effect (Cohen’s *f* ≥ .4).

Screening ensured that subjects were right-handed, learned English as their first language, had normal/corrected vision, and had no history of neurological disorder. Informed consent was obtained in accordance with the MU Institutional Review Board. Seven subjects were excluded from all analyses due to not attending the second session (3 subjects), excessive artifact in the EEG (2 subjects), behavioral performance that was >2 SDs below the group mean in a condition of interest (1 subject), or technical errors (1 subject). The remaining 17 subjects (6 females, 11 males) were 18-22 years old (M = 19).

### 2.2 Stimuli and design

Stimuli were 400 color pictures of common, nameable objects (cf. the words in Johnson & McGhee, 2015). For each subject, two sets of 100 pictures each were randomly selected to form the remote and recent encoding lists. Within each encoding list, 50 of the pictures appeared only once (hereafter, *1x*), and the remaining 50 pictures appeared four times each (*4x*), resulting in a total of 250 trials per list. Four targeted test lists were then constructed for each subject by randomly selecting 30 *target* items (out of the 50 possible) from the corresponding week (remote/recent) × repetition (1x/4x) condition.

Additionally, 20 items were selected from the encoding lists corresponding to the alternative week (e.g., when the remote condition was targeted, these items were drawn equally from the *recent-1x* and *recent-4x* conditions) to serve as *nontargets*. This selection approach for nontargets encouraged subjects to focus their orienting strategy exclusively on a particular week, which corresponded to our primary interest, as opposed to orienting to a specific week × repetition condition (as would be the case if we selected nontargets from all of the remaining conditions). A further 30 *new* items were selected from the stimulus pool for each targeted list. A final recognition test list comprised the 120 new items from across all of the targeted tests along with the 80 remaining items from the pool.

All pictures were displayed on a solid gray square at the center of the black background of a 24-inch widescreen LCD monitor (cropped to 1024×768 resolution). At the 1 m viewing distance, pictures subtended approximate visual angles of 3.1° and the gray square subtended 4.6°. All instructions, response options, and fixation crosses were displayed in white Arial font. The Cogent 2000 toolbox (http://www.vislab.ucl.ac.uk) was used to control stimulus presentation in MATLAB (The MathWorks, Natick, MA).

### 2.3 Procedure

Subjects completed the remote encoding phase during the first visit to the laboratory and then returned approximately one week (range = 6-8 days) later to complete the recent encoding phase, targeted memory test phase, and final recognition memory test. The first and second visits respectively lasted about 30 minutes and two hours.

For each encoding phase, subjects made pleasantness ratings for a series of pictures by using their right index through little fingers to respectively press keyboard keys for “very pleasant”, “somewhat pleasant”, “somewhat unpleasant”, and “very unpleasant”. Subjects were informed that some pictures would occasionally repeat, and it was emphasized that responses should be based on how they felt in that moment rather than thinking back to previous presentations. Pictures presented either once or four times were randomly intermixed. For the recent encoding phase, subjects were additionally informed that none of the pictures had been presented during the previous (remote) list. Each picture was presented for 2 s and followed by a central fixation cross for .5 s. Subjects were instructed to try to respond before the next picture appeared. A short break occurred at the midpoint of each list.

The recent encoding phase was followed by EEG preparation, lasting about 30 minutes, and then the targeted memory test phase. To minimize any differences in strategies for the two encoding phases, subjects were kept unaware of the nature of the memory test until immediately before it began. The targeted test phase was divided into four blocks, each of which employed an exclusion procedure (Jacoby, 1991) to target pictures from one of the four study conditions (*remote-1x*, *remote-4x*, *recent-1x*, and *recent-4x*). For example, in the block targeting remote-1x pictures, those items required a keypress response made with the right index finger, whereas items encoded in the recent list (nontargets) and new items both required a keypress with the right middle finger. To help prevent the exclusion task from being too challenging, nontargets always came from the alternative encoding list rather than from the alternative repetition condition in the targeted list, and the instructions emphasized rejecting items from the other encoding list. Each test picture was presented for 3 s and followed by a central fixation cross for 1 s. Subjects were instructed to respond before the next picture appeared. Test blocks were separated by breaks of about five minutes each, during which instructions and practice on the upcoming block were provided. Blocks targeting the remote and recent lists followed an ABBA order that was counterbalanced across subjects, whereas targeting the 1x/4x conditions was randomized.

Finally, subjects undertook a recognition test that assessed memory for new items presented during the targeted test phase. Such memory tests—sometimes called *memory-for-foils* tests (Jacoby et al., 2005; Shimizu & Jacoby, 2005)—have been used in other contexts to examine the subsequent behavioral consequences of adopting different retrieval orientations (also see Johnson & McGhee, 2015). For the final test, subjects were instructed to make one keypress to pictures presented during any targeted test block and an alternative keypress to pictures that were entirely new to the experiment (using the right index and middle fingers, respectively). Trial timing was the same as in the targeted test, and a short break occurred at the midpoint of the test.

### 2.4 EEG acquisition and processing

EEG data were continuously recorded during the targeted test phase from 59 Ag/AgCl sintered ring electrodes embedded in an elastic cap (Easycap, Herrsching, Germany; http://www.easycap.de), using a BrainAmp Standard system (BrainVision LLC; Durham, NC; http://www.brainvision.com). Electrode placement was based on the extended 10-20 system and included: Fpz/1/2, AFz/3/4/7/8, Fz/1-8, FC1-6, FT7/8, Cz/1-6, T7/8, CPz/1-6, TP7/8, Pz/1-8, POz/3/4/7/8, and O1/2. Additional electrodes were used for the online reference (at FCz), ground (FT10), EOG (IO1 and LO1/2), and offline re-referencing (M1/2). Impedances were adjusted below 5 kΩ before recording started, and the data were recorded at a sampling rate of 1 kHz and amplifier bandwidth of .01-100 Hz.

Offline processing of the EEG data was implemented with the EEGLAB toolbox (Delorme & Makeig, 2004; https://sccn.ucsd.edu/eeglab/). The data were merged across test blocks, re-referenced to linked mastoids, downsampled (200 Hz), bandpass filtered (.05-20 Hz), epoched from -0.2 s to 2 s relative to each test item onset, and baseline-corrected across the pre-stimulus period. Independent component analysis (ICA) was used to identify components corresponding to blinks and eye-movements, which were rejected based on high correlation (>3 SDs) with the bipolar EOG (Fp1-IO1 and LO1-LO2) time courses. Epochs containing considerable artifact were next manually rejected before completing a second pass of ICA. Components reflecting artifact were manually rejected based on their scalp topography and power spectra. The FASTER toolbox (Nolan et al., 2010) was then used to identify bad channels on an epoch-wise basis, with thresholds of ≥3 SDs employed for cross-channel correlation, variance, and the Hurst exponent. Finally, bad channel epochs were interpolated using a spherical spline function. The across-subject mean numbers of epochs forming the new-item ERPs for each condition were: remote-1x = 24.5 (range: 19-30), remote-4x = 27.9 (25-30), recent-1x = 28.6 (22-30), and recent-4x = 28.3 (26-30).

### 2.5 ERP analysis

In addition to conducting pre-experimentally planned ERP analyses using an electrode montage (see Figure 1) and latency intervals that have been previously employed to identify retrieval orientation effects (e.g., Dzulkifli & Wilding, 2005; Johnson & McGhee, 2015; Johnson & Rugg, 2006; Robb & Rugg, 2002), permutation-based approaches were used to further test and characterize the spatial and temporal distributions of the effects (Maris & Oostenveld, 2007). Analysis in the spatial domain used ERP amplitude averages at each electrode for each of the foregoing latency intervals, and electrodes were defined as neighbors based on an adjacency template (Easycap M11) in the Fieldtrip toolbox (Oostenveld et al., 2011; http://fieldtrip.fcdonders.nl). For the temporal domain, electrode-wise averages were subdivided into 100-ms intervals (0-100 to 1900-2000 ms), with electrode and interval adjacency contributing to contiguous clusters. Both approaches involved randomly shuffling the condition labels for each subject 1000 times. The maximum number of clustered electrodes or clustered electrode × interval elements that were significant, based on paired *t*-tests (*p* < .025 in each direction), were identified for each permutation. (The results were similar for the two directions, so we conservatively applied the largest cluster size for both cases.) The actual (non-shuffled) results were then compared against the permuted values to determine cluster-corrected significance.

**Figure 1:**
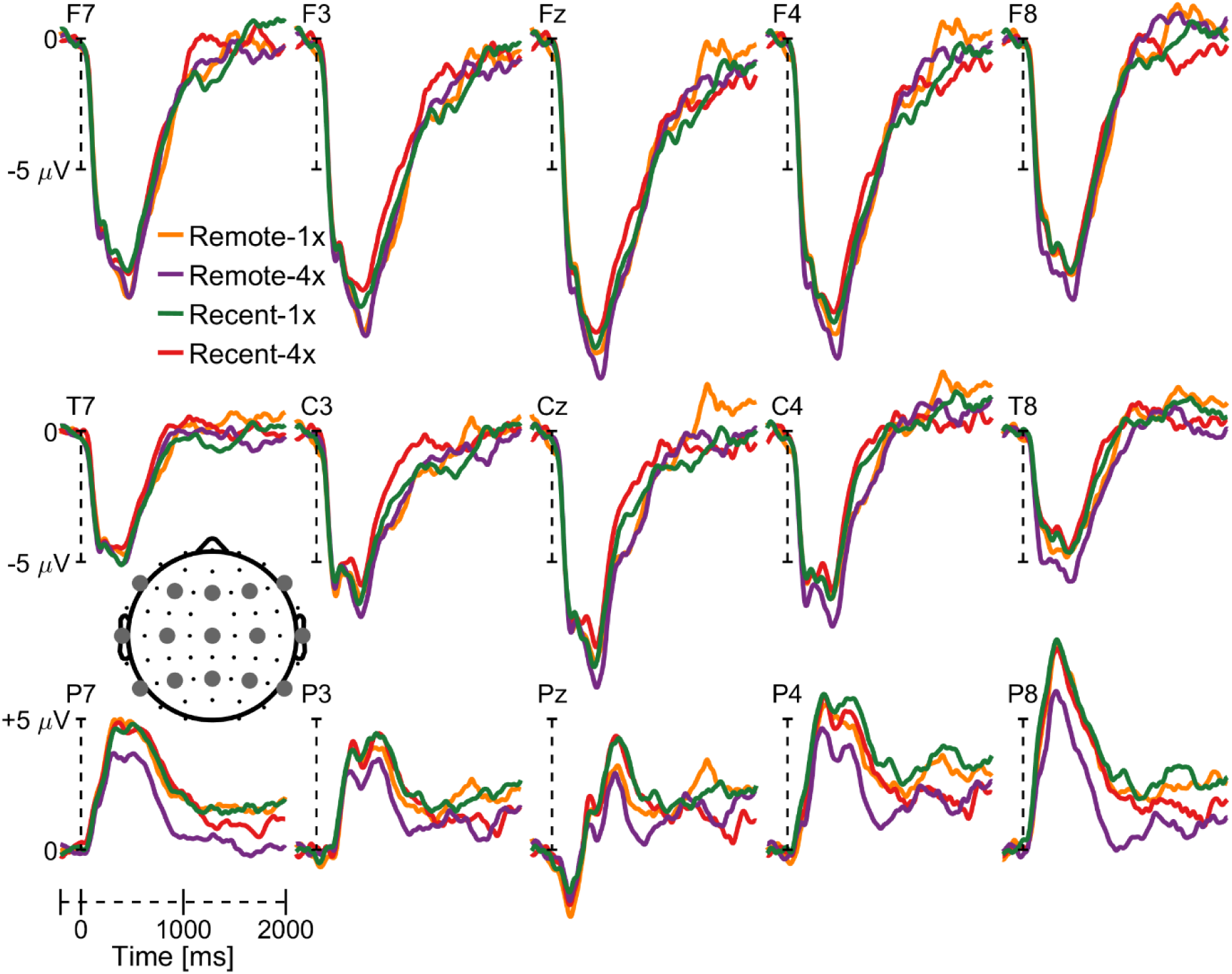
ERPs corresponding to correctly-rejected new items. Grand-average new-item ERPs are provided for the four types of targeted memory tests. Data are provided from a 15-electrode montage (see inset) that also corresponds to the main analyses reported in the text. These ERPs were smoothed using a 100-ms moving average for display purposes only. See the online version for color.

## 3. RESULTS

### 3.1 Behavioral results

Correct response proportions and response times (RTs) from the targeted test phase are summarized in Table 1. These measures were separately submitted to three-way ANOVAs, including factors of item type (target/nontarget/new), week (remote/recent), and repetition (1x/4x; *df*s were Greenhouse-Geisser adjusted when appropriate). The ANOVA of proportions gave rise to significant main effects of item type (*F*_1.1,17.9_ = 27.05, *p* < .001), week (*F*_1,16_ = 72.21, *p* < .001), and repetition (*F*_1,16_ = 86.60, *p* < .001), and significant interactions of item type × week (*F*_1.4,21.7_ = 29.29, *p* < .001), item type × repetition (*F*_1.4,23_ = 9.40, *p* = .002), week × repetition (*F*_1,16_ = 25.73, *p* < .001), and item type × week × repetition (*F*_1.7,27_ = 9.99, *p* = .001). For the correct RTs, ANOVA revealed all significant main effects (item type: *F*_1.4,21.7_ = 30.40, *p* < .001; week: *F*_1,16_ = 19.08, *p* < .001; repetition: *F*_1,16_ = 11.97, *p* = .003) and an item type × week interaction (*F*_1.9,30.7_ = 18.08, *p* < .001).

**Table 1:** Behavioral results for the targeted memory tests. Mean proportions of correct responses and associated response times (RTs) for the targeted memory test phase

|  | Targeted condition |  |  |  |
| --- | --- | --- | --- | --- |
| Item type | Remote-1x | Remote-4x | Recent-1x | Recent-4x |
| *Proportions* |  |  |  |  |
| **Target** | .52 (.06) | .73 (.05) | .83 (.04) | .93 (.02) |
| **Nontarget** | .94 (.02) | .97 (.01) | .89 (.02) | .97 (.01) |
| **New** | .85 (.02) | .95 (.01) | .97 (.02) | .98 (.01) |
| *RTs* |  |  |  |  |
| **Target** | 1459 (79) | 1305 (60) | 1215 (49) | 1104 (42) |
| **Nontarget** | 1144 (74) | 1104 (50) | 1121 (54) | 1083 (56) |
| **New** | 1231 (84) | 1089 (60) | 1065 (51) | 971 (51) |

SEMs in parentheses; RTs are in milliseconds.

To further understand the above interactions, and because we eventually focus on correct rejections for the ERP analyses, follow-up ANOVAs were restricted to the new items. ANOVA of the proportions revealed significant main effects (week: *F*_1,16_ = 37.77, *p* < .001; repetition: *F*_1,16_ = 12.92, *p* = .002) and an interaction (*F*_1,16_ = 34.70, *p* < .001). The interaction can be described as additional repetitions of the targeted condition increasing the rejection rate for the remote condition (.85 vs. .95; *t*_16_ = 5.00, *p* < .001), whereas the corresponding rates for the recent condition were already near perfect (.97 vs. .98; *t*_16_ = .92, *p* = .37). ANOVA of the new-item RTs gave rise only to significant main effects (week: *F*_1,16_ = 14.69, *p* = .001; repetition: *F*_1,16_ = 13.91, *p* = .002), reflecting faster RTs for the recent and repeated (4x) conditions. Given the motivation to equate performance across the remote and recent tests, we also directly contrasted the data for the remote-4x and recent-1x conditions, revealing no differences for either measure (proportions: *t*_16_ = .59, *p* = .56; RTs: *t*_16_ = .70, *p* = .49).

Finally, to evaluate the consequences of targeting different encoding conditions on subsequent performance, the behavioral measures for the memory-for-foils (see Table 2) were analyzed. As our main interest was in new items (i.e. foils) from the previous targeted tests, ANOVAs included factors of week (remote/recent) and repetition (1x/4x). No significant effects were evident for correct proportions (*p*s > .19) or RTs (*p*s > .34). Pairwise comparisons of foils from the remote-4x and recent-1x tests also indicated no reliable differences (proportions: *t*_16_ = 1.50, *p* = .15; RTs: *t*_16_ = .79, *p* = .44).

**Table 2:**
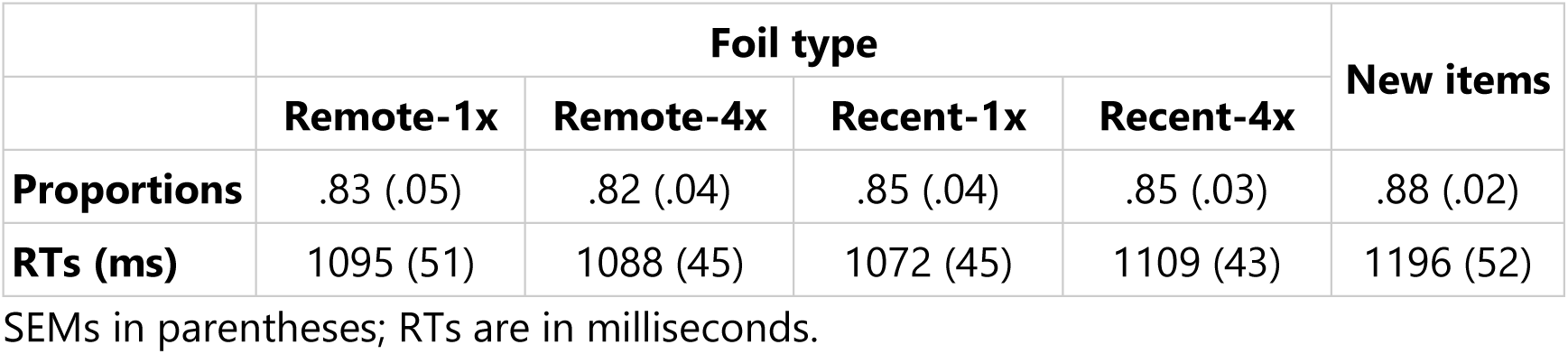
Behavioral results for the memory-for-foils test. Mean proportions of correct responses and associated response times (RTs) for the memory-for-foils test

|  | Foil type |  |  |  | New items |
| --- | --- | --- | --- | --- | --- |
|  | Remote-1x | Remote-4x | Recent-1x | Recent-4x |  |
| **Proportions** | .83 (.05) | .82 (.04) | .85 (.04) | .85 (.03) | .88 (.02) |
| **RTs (ms)** | 1095 (51) | 1088 (45) | 1072 (45) | 1109 (43) | 1196 (52) |

SEMs in parentheses; RTs are in milliseconds.

### 3.2 ERP results

The ERPs corresponding to correctly rejected new items were initially contrasted across the remote-4x and recent-1x conditions to identify retrieval orientation effects while minimizing behavioral differences (i.e. difficulty). ERPs associated with these and the remaining new-item conditions are shown in Figure 1. As shown, beginning at about 500 ms after stimulus onset, the waveforms over posterior scalp appear more positive for the recent-1x compared to remote-4x condition. The mean ERP amplitudes for the two conditions were extracted from each electrode in the displayed montage during six latency intervals: 200-500, 500-800, 800-1100, 1100-1400, 1400-1700, and 1700-2000 ms. ANOVA employing factors of orienting condition (remote-4x/recent-1x), anterior/posterior chain (frontal/central/parietal), laterality (inferior left/superior left/midline/superior right/inferior right), and interval gave rise to a significant orienting condition × anterior/posterior chain interaction (*F*_1.7,27.1_ = 7.36, *p* = .004). No other effects involving the orienting factor were significant (*p*s > .08).

To better assess the onset and time course of the orienting effects, planned ANOVAs were conducted for each latency interval (employing the remaining factors; for similar approach, see Dzulkifli & Wilding, 2005; Johnson & McGhee, 2015; Johnson & Rugg, 2006; Robb & Rugg, 2002). These analyses gave rise to significant orienting condition × anterior/posterior chain interactions for the 500-800 (*F*_1.3,20.7_ = 4.18, *p* = .045), 800-1100 (*F*_1.5,24.7_ = 9.15, *p* = .002), 1100-1400 (*F*_1.5,23.2_ = 7.83, *p* = .005), and 1400-1700 ms (*F*_1.6,26.2_ = 8.26, *p* = .003) intervals. Figure 2A highlights the fact that the orienting differences were restricted to posterior scalp, and follow-up ANOVAs at each anterior/posterior chain confirmed significant orienting main effects across parietal electrodes at 500-800 (*F*_1,16_ = 6.48, *p* = .022), 800-1100 (*F*_1,16_ = 5.84, *p* = .028), and 1400-1700 ms (*F*_1,16_ = 6.36, *p* = .023). Similar analyses focused on the frontal and central electrodes in these three intervals gave rise to no significant effects (*p*s > .26). For the 1100-1400 ms interval, testing each electrode chain revealed no significant effects (*p*s > .21).

**Figure 2:**
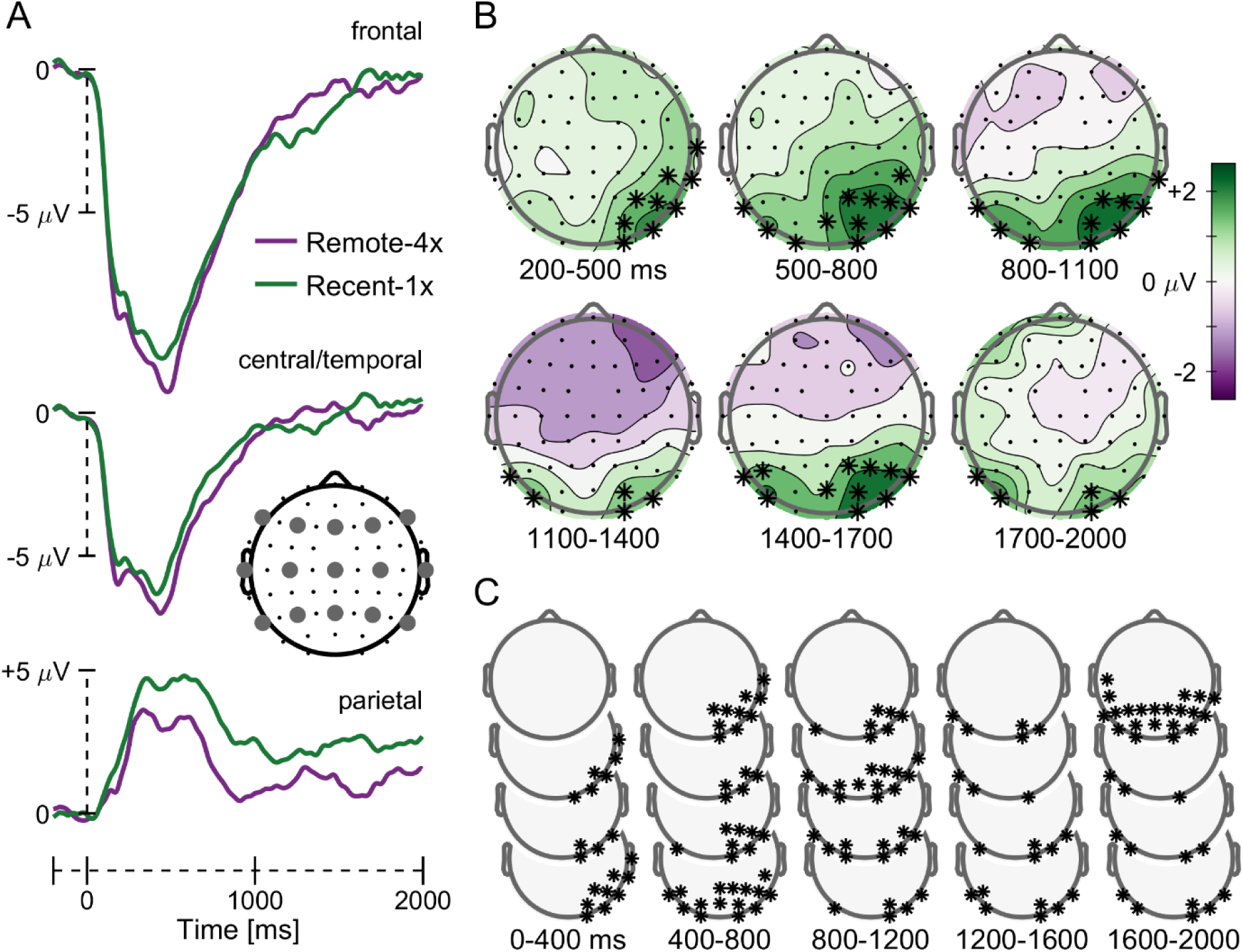
ERP results of orienting retrieval to memory age. (A) Grand-average ERPs collapsed across three anterior/posterior electrode chains (see inset for the montage) according to whether retrieval was oriented toward items presented four times one week ago (remote-4x) or one time 30 minutes ago (recent-1x). (B) Topographic scalp distribution of the recent-1x minus remote-4x orienting differences for new items in each latency interval used for the main analyses. Asterisks indicate significant differences (*p* < .05, uncorrected), with the right-lateralized clusters during the 200-500, 500-800, and 1400-1700 ms intervals passing the cluster-corrected threshold. (C) Scalp distribution of significant orienting differences during 100-ms latency intervals throughout the recording epoch. Asterisks indicate a single cluster of electrodes that passed the cluster-corrected level of significance based on time and topography. See the online version for color.

The broader topographic distribution of the orienting differences was further tested with permutation analyses that controlled the FWER across clustered electrodes. These analyses were conducted in two ways. First, the electrode-wise differences between the recent-1x and remote-4x conditions were averaged within the six previously-defined latency intervals. (We conducted permutations within every interval, irrespective of the significance of effects reported above.) The resulting maps are displayed in Figure 2B and revealed significant clusters of electrodes (passing the minimum size of 8) over right posterior scalp in the 200-500, 500-800, and 1400-1700 ms intervals. The second permutation analysis, based on combining adjacent electrodes and consecutive latency intervals (100-ms each), was used to better assess the time course of the orienting effects. As shown in Figure 2C, the posterior orienting effect was more sustained than the foregoing analysis indicated, beginning in the 100-200 ms bin and lasting throughout the recording epoch. This effect is linked into a single significant cluster of 145 electrode × time interval elements (vs. critical size of 103).

Although our repetition manipulation provided conditions where the remote and recent conditions could be matched on behavior, our final analysis sought to dissociate the neural correlates of difficulty and orienting. We focused on the repetition effect within the remote condition, as it was accompanied by significant behavioral differences (see Table 1). ANOVA of the new-item ERPs from tests targeting remote-4x and remote-1x items (see Figure 1) included the latency interval and electrode factors previously used, along with a factor accounting for repetition condition. Although the omnibus ANOVA revealed no significant effects involving the repetition factor (*p*s > .21), planned ANOVAs within each interval gave rise to a significant repetition × anterior/posterior chain × laterality interaction for 200-500 ms (*F*_3.9,61.9_ = 3.31, *p* = .017; no other latencies were associated with significant effects, all *p*s > .08). In this early interval, pair-wise *t*-tests at each of the 15 electrodes in the montage revealed that repetition differences were restricted to the P7 and P8 sites (respectively, *t*_16_ = 2.65 and 3.35, *p* = .018 and .004; *p*s > .09 for the other electrodes). Figure 3A shows the effects at these sites in more detail, whereby the remote-1x waveform was more positive than that for remote-4x.

**Figure 3:**
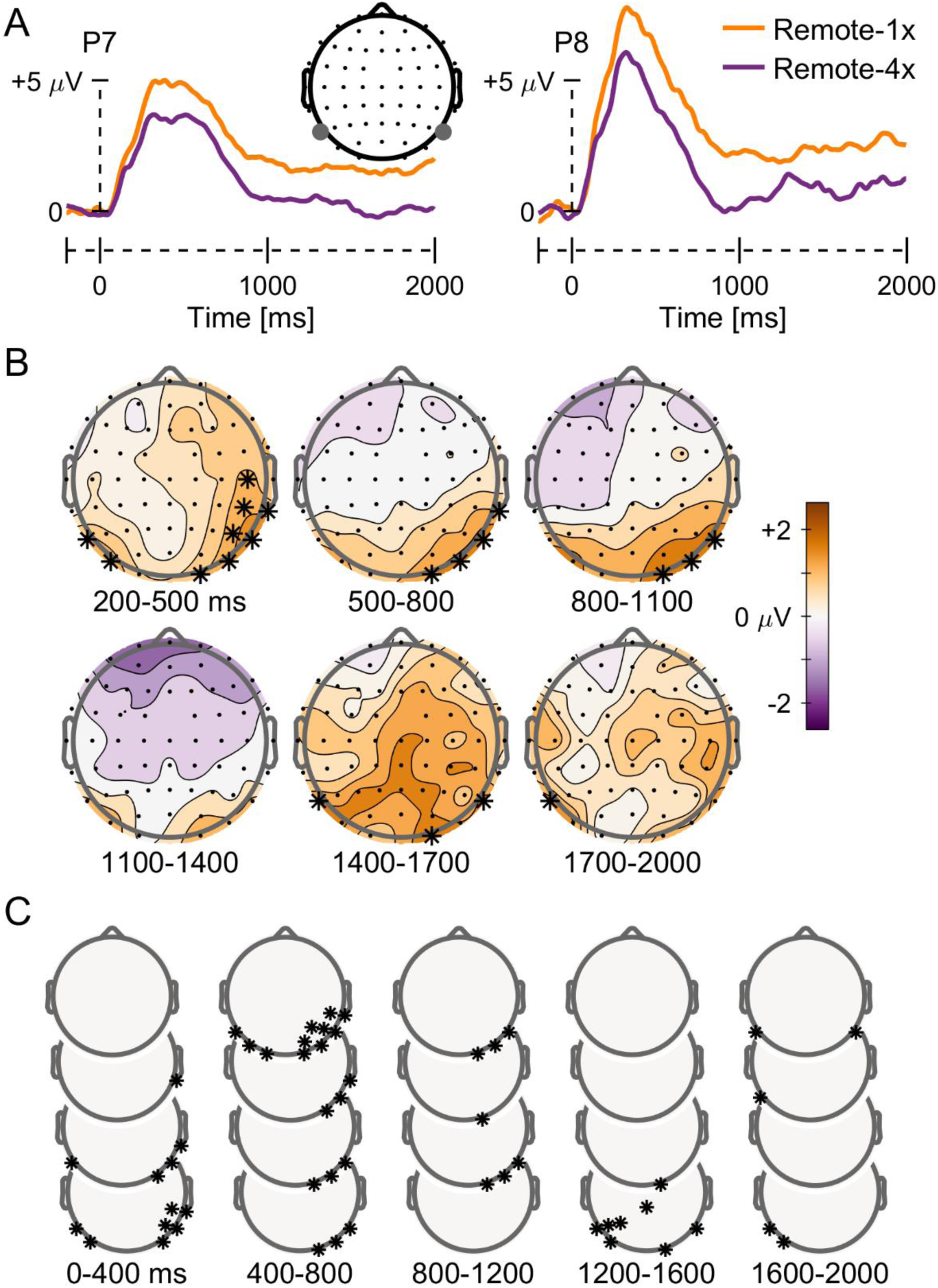
ERP results of retrieval difficulty. (A) Grand-average ERPs from two electrodes (P7 and P8) exhibiting repetition-related differences for new items when targeting items from one week ago (i.e. remote-4x vs. remote-1x). (B) Topographic scalp distribution of the remote-1x minus remote-4x differences for new items, with asterisks indicating significant electrode-wise effects (*p* < .05, uncorrected; none of these effects met the cluster-corrected threshold). (C) Topographic scalp distribution of the significant (uncorrected) new-item repetition differences for 100-ms latency intervals, which also failed to pass cluster correction. See the online version for color.

Turning to the topography of the repetition effect, as displayed in Figure 3B, we next sought to better assess the extent and timing of this effect. Permutations within the predetermined latencies revealed two separate clusters of 7 and 2 electrodes during the 200-500 ms interval, although the larger of these did not pass the critical threshold of 8 electrodes. Additionally, the largest cluster in subsequent intervals only comprised 4 electrodes. For the electrode × time (100-ms bins) permutations, the evidence of any effects over posterior scalp was still insignificant (see Figure 3C). In particular, an effect over right posterior sites lasted from the 100-200 to 1000-1100 ms intervals, but it comprised only 33 clustered elements (compared to the critical threshold of 101). Clusters over left posterior scalp only reached a maximum size of 6 data points.

## 4. DISCUSSION

The experiment reported here investigated the differential engagement of retrieval processing when remote and recent memories are targeted. Specifically, we examined the effects of orienting retrieval to stimuli encoded one week versus about 30 minutes earlier, and whether the ERP correlates of orienting were distinct from those related to the increased difficulty associated with accessing remote memories. To this end, we directly contrasted strategic orienting to remote target memories that had been studied repeatedly (four times) and recent memories that were studied only once, resulting in equivalent behavioral performance in the form of both accuracy and RTs for new test items. Consistent with our previous investigation of retrieval orientation for memory age (Johnson & McGhee, 2015), the ERPs elicited by correctly-rejected new items here were more positive-going when targeting recent compared to remote memories, and these effects were evident by about 500 ms post-stimulus and maximal over posterior scalp.

The repetition manipulation employed in the current study also allowed for distinguishing ERP differences putatively related to retrieval difficulty from those associated with strategic orienting toward memory age. This distinction was apparent in both the timing and the polarity of the ERP effects. While the repetition effect (remote-4x vs. remote-1x) appeared early (200-500 ms post-stimulus onset) and was short-lived, the orienting effect onset later (500-800 ms) and lasted until about the end of the recording epoch. Moreover, the repetition effect took the form of positive-going waveforms for the more difficult condition (i.e. remote-1x > remote-4x; for analogous results, see Robb & Rugg, 2002), whereas the opposite pattern emerged for orienting, with the condition that would presumably be “easier” according only to memory age (i.e. irrespective of repetition) eliciting relative positivity (i.e. recent-1x > remote-4x). That is, though the concern motivating this study was that ERP correlates of difficulty could be posing as those related to differential orienting to memory age, the present pattern of results suggests that difficulty could have actually reduced the magnitude of the orienting differences observed.

The argument about difficulty stated above could also be applied to the findings of our previous study that investigated differential orienting to memory age (Johnson & McGhee, 2015). In that study, although we attempted to control for difficulty differences in a post hoc manner, some residual – and potentially undetected – differences in the new-item accuracy rates could have influenced the orientation effects. Similarly, for the current study, although we took care in matching behavioral performance (via a series of pilot behavioral experiments that tested different materials and levels of repetition) for the new test items that were of interest, other remaining differences are more difficult to control. These uncontrolled differences can involve items such as targets and nontargets, as is apparent in Table 1 (compare the Remote-4x and Recent-1x columns). For this and other reasons noted in the Introduction, these items were not included in our ERP analyses. It would be naïve, though, to assume that the traditional behavioral measures we employed here (proportion correct and RT) are exhaustive in their ability to detect differences in behavior, let alone the underlying cognitive or physiological processes. Additionally, other issues such as floor and ceiling effects are potentially overlaid on our measurement abilities. Nonetheless, we think the current approach of attempting to isolate effects of difficulty and then dissociating those from the effects of interest (for analogous approaches, see Dzulkifli et al., 2004; Herron & Wilding, 2004; Robb & Rugg, 2002) has utility in further understanding retrieval strategies.

Although it was not relevant to the main ERP findings from the targeted memory phase, another noteworthy aspect of our results is the lack of behavioral differences on the final memory-for-foils test. This null finding contrasts with the findings several recent studies (e.g., Danckert et al., 2011; Jacoby et al., 2005; Marsh et al., 2009; Shimizu & Jacoby, 2005; Vogelsang et al., 2016; but for other null results, see Johnson & McGhee, 2015, and Kantner & Lindsay, 2013). For example, in Vogelsang et al. (2016), new test items were more likely to be subsequently retrieved on a later test if they were presented in the context of a memory task targeting deeply versus shallowly encoded memories. The discrepancy between the current and previous results could be due, on one hand, to the nature of the task manipulations employed during the retrieval phase, such that the levels-of-processing (deep vs. shallow) manipulation in studies such as Vogelsang et al. (2016) was more powerful than the remote-recent targeting task we employed. Alternatively, the pictures used in our study gave rise to better memory performance overall on the final test, compared to that for the words employed by Vogelsang et al. (2016), thus potentially restricting the range available for observing any difference. Finally, a difference in statistical power between that study and ours also potentially accounts for the discrepancy; a *post hoc* power analysis based on the effect size (d_z_ = .75) in Vogelsang et al. (2016) indicated 80% power with 16 subjects. Further studies directed at bridging the disparate findings from these different types of manipulations in terms of behavioral consequences on memory are clearly needed.

To summarize, the current study extends previous investigations of retrieval orientation strategies toward memory age (also see Johnson & McGhee, 2015; Roberts et al., 2014) by showing that the ERP correlates of such orienting are largely dissociable from those related to retrieval difficulty. It remains to be determined what characteristics of memory traces might be driving the differential adoption of orienting processes at the time of retrieval. As we have previously speculated (Johnson & McGhee, 2015), it is possible that these orienting effects reflect the setting of expectations about differences in the quality or amount of detail (Vilberg & Rugg, 2009) that should accompany successful retrieval depending on the recency of encoding. Likewise, the effects could signify the reinstatement of self-referential or other temporally-distinct contextual information that potentially serves the purpose of matching against the products of retrieval. The present study cannot distinguish between these possible mechanisms. However, considering that the orienting effects remained when behavioral performance was matched, the current results appear inconsistent with the notion that subjects are focusing cue processing on the relative strength of the targeted memories and the corresponding difficulty (or effort) associated with overcoming that quality (cf. Johnson & McGhee, 2015). Nevertheless, these findings provide further support that the ages of different memories alone, or perhaps the temporal contexts associated with them, can be used as the basis for directing cognitive control processes at the time of episodic retrieval, presumably in service of maximizing retrieval success.

## ACKNOWLEDGEMENTS

We thank Bruce Bartholow, Roxana Botezatu, and Steve Hackley for helpful feedback on an earlier version of this article. The authors report no conflicts of interest.

